# Identification of genes affecting saturated fat acid content in *Elaeis guineensis* by genome-wide association analysis

**DOI:** 10.1101/341347

**Authors:** Wei Xia, Tingting Luo, Wei Zhang, Annaliese S. Mason, Dongyi Huang, Xiaolong Huang, Wenqi Tang, Yajing Dou, Chunyu Zhang, Yong Xiao

## Abstract

Oil palm is the highest yielding oil crop per unit area worldwide. Unfortunately, palm oil is often considered unhealthy. In particular, palmic acid (C16:0) is a major component of palm oil. In this study a total of 1 261 501 SNP markers were produced in a diversity panel of 200 oil palm individuals. Oil content in this population varied from 29.8% to 70.3%, palmic acid varied from 31.3% to 48.8%, and oleic acid varied from 31.3% to 50.1%. We identified 274 SNP markers significantly associated with fatty acid compositions; 44 candidate genes in the flanking regions of these SNPs were involved in fatty acid biosynthesis and metabolism. Among them, two acyl-ACP thioesterase B genes had differential expression patterns between the mesocarp and kernel, tissues which show different oil profiles in oil palm (high palmic acid and high lauric acid respectively). Overexpression of both genes caused a significant increase in palmic acid content, while overexpression of the *EgFatB2* gene also caused an accumulation of lauric acid and myristic acid. Our research provides genome-wide SNPs, a set of markers significantly associated with fatty acid content, and validated candidate genes for future targeted breeding of lower saturated fat content in palm oil.

## Introduction

Oil palm (*Elaeis guineensis*, 2n = 32), belongs to the palm family (Arecaceae), and is an important tropical oil crop grown throughout Southeast Asia, Africa, Central America and Brazil. Oil palm has the highest oil yield per unit area of all oil crops: more than six times that of peanut, eight times that of soybean, and ten times that of rapeseed (Huang *et al.,* 2017). The world production of palm oil in 2017 was approximately 65 million tonnes (http://faostat3.fao.org/home/E). The two oil storage tissues of oil palm are the mesocarp and kernel, each of which produces oil with different fatty acid compositions. Palm oil usually refers to the oil extracted from the oil palm mesocarp, where palmic acid (16:0) is a dominant fatty acid (50%), while in kernel oil, lauric acid (12:0) is the major fatty acid (50%). In oil palm, one of the principal breeding objectives is to decrease palmitic acid content (16:0) and increase oleic acid (18:1) and linoleic acid content (18:2) in palm oil. Recently, breeders in Malaysia have obtained an oil palm variety with high oleic acid (approximately 52%), although this is still far lower than the oleic acid content in peanut (more than 75% in high oleic acid varieties) and rapeseed (more than 80%) (Rajanaidu *et al.,* 2017). However, applying conventional breeding processes to improve fatty acid composition is slow in oil palm because of the long life cycle of the plant. Identification of the underlying genes controlling oil composition and subsequent production of molecular markers linked to low palmitic acid content would be extremely helpful in speeding up breeding and selection processes in oil palm.

In *Arabidopsis*, a great deal of work has been done to elucidate genes involved in the fatty acid biosynthesis pathway, which occurs mainly in the plastids and endoplasmic reticulum. Initially, acetyl-CoA as precursor is converted into Malonyl-CoA catalyzed by acetyl CoA carboxylase (ACCase) (Sasaki *et al.,* 1997). Malonyl-CoA is subsequently polymerized at a frequency of two carbons per cycle into the Acyl carbon chain and combined with acyl carrier protein (ACP) catalyzed by ketoly-ACP synthase (KAS) (Choi *et al.,* 2000). Finally, the elongation of the carbon chain is terminated catalyzed by acyl-CoA thioesterase (FAT) (Hunt *et al.,* 2002). Previous research has showed that acyl-CoA thioesterase (FAT) can be divided into FATA and FATB (Salas *et al.,* 2002). FATA was thought to be able to bring about termination of C18:0-ACP and C18:1-ACP (Mekhedov *et al.,* 2000). However, FATB may also be involved in termination of saturated fatty acyl-ACP (Bonaventure *et al.,* 2004). In *Arabidopsis thaliana*, overexpressing the *AtFatB1* gene can significantly increase palmic acid content. The disruption of FatB expression resulted in a 56% reduction in palmitic acid and a 50% reduction of stearate in *Arabidopsis* seeds (Wilson *et al.,* 2001; Buhr *et al.,* 2002). Moreover, enhanced expression of FatB from *Umbellularia californica* can significantly increase lauric acid content (C12:0) in *Brassica napus* (Voelker *et al.,* 1996).

Association mapping is an efficient method to elucidate the genetic bases of complex agronomic traits and to identify molecular markers associated with agronomic traits based on natural populations with or without pedigree relationships (Flint-Garcia *et al.,* 2003). The resolution of an association study depends on the extent and structure of linkage disequilibrium (LD) in the selected population. With the development of sequencing technology and high-throughput single nucleotide polymorphism (SNP) markers, association mapping has been widely applied in different crops to dissect complex agronomic traits (Atwell *et al.,* 2010; Li *et al.,* 2013; Huang *et al.,* 2011). However, there is still the problem of how to reduce the detection of spurious associations between traits and markers resulting from population structure. One solution is to perform association analysis between allelic and phenotype variation in a less structured population (Pritchard *et al.,* 2000). Another solution for structured populations is to apply a mixed-model approach, which can decrease spurious associations (Sorkheh *et al.,* 2008). Compared with QTL (quantitative trait loci) mapping in biparental cross populations, association analysis using natural populations is more time- and cost-effective, especially for woody perennial crops with long life cycles or which require large planting areas. Previously in oil palm, efforts have been made to identify QTL for agronomic traits for marker-assisted selection based on biparental cross populations (Rance *et al.,* 2001; Billotte *et al.,* 2010; Jeennor *et al.,* 2014; Tisne *et al.,* 2015). QTL mapping has been used to resolve the genetic bases of several complex traits in oil palm, including yield components (Rance *et al.,* 2001; Billotte *et al.,* 2010), fatty acid composition (Singh *et al.,* 2009; Montoya *et al.,* 2013), sex ratio (Ukoskit *et al.,* 2014) and embryogenesis (Ting *et al.,* 2013). However, the contribution of QTL mapping studies to marker-assisted breeding outcomes has been less than expected. Although association mapping has been validated to be a reliable method for identifying trait-associated markers for marker-assisted selection, this method has rarely been applied in oil palm.

High-density molecular markers are a prerequisite for genome-wide association analysis (GWAS) (Kenny *et al.,* 2011). Specific locus amplified fragment sequencing (SLAF-seq) is a fast, accurate, highly efficient, and cost-effective method for developing high-density SNP and Indel markers (Sun *et al.,* 2013; Zhang *et al.,* 2013). In the present study, we developed genome-wide SNP markers using SLAF-seq technology from oil palm 200 individuals with different geographical origins, and used this data to evaluate the genetic diversity population structure and patterns of linkage disequilibrium in this population. Subsequently, we,investigated fatty acid content and identified SNP markers linked with this trait by GWAS analysis, identifying candidate genes adjacent to these SNP markers with known roles in fatty acid content and analyzing their expression patterns in different tissue types. To validate the role of top candidate genes in regulating palmic oil content, we transferred these genes into *Arabidopsis thaliana*. Our study provides a comprehensive understanding of fatty acid content in oil palm, and SNPs and candidate genes detected will play an important role in breeding for fatty acid content in oil palm. Finally, we discuss how best to utilize our results to decrease palmic acid content (C16:0) and increase oleic acid content (18:1) in oil palm, producing a healthier oil for human consumption.

## Materials and methods

### Plant materials and DNA extraction

A total of 200 oil palm individuals were collected from different geographical districts: 21 were collected from southern China (Hainan province), which were introduced from Malaysia to China in the 1960s; 80 were collected from the plateau region of Malaysia; 90 were collected from Costa Rica; and the remaining nine individuals were collected from Africa. The 200 oil palm individuals were still grown in oil palm germplam resources of Coconut Research Institute of Chinese Academy of Tropical Agricultural Sciences in Wenchang town of Hainan province of China since 2008. The average temperature and humidity are approximately 23.9 ^0^C and 89%. DNA samples were prepared from young leaves using the mini-CTAB method (Murray and Thompson, 1980). The concentration and quality of the 200 oil palm DNA samples was examined using a Nanodrop 2000 UV-Vis spectrophotometer (NanoDrop, Wilmington, DE, USA). The quantified DNA was diluted to 100 ng/μl for SLAF sequencing.

### Selection of enzyme combinations

In order to obtain more SLAF tags, *in silico* restriction enzyme cutting sites across the oil palm genome were analyzed to select restriction enzyme combinations matching the following criteria: (1) resulting SLAF tags should contain a low percentage of repeat sequences; (2) SLAF tags should be evenly distributed across the genome of *Elaeis guineensis*; (3) simulated fragments must align uniquely to the reference genome; (4) larger number of SLAF tags predicted (the length of SLAF tags varied from 314 to 414 bp). According to these four criteria, the restriction enzyme combination of HaeIII + Hpy166II was selected for subsequent experiments. A total of 244 702 SLAF tags were predicted based on *in silico* digestion using the restriction enzyme combination of HaeIII + Hpy166II.

### SLAF sequencing

Genomic DNA extracted from the spear leaves of each oil palm accession was digested with HaeIII and Hpy166II to get SLAF tags. Subsequently, the obtained SLAF tags were used as DNA templates for fragmentation and reparation, dual-index paired-end adapter ligation, PCR amplification, and target fragment selection (Sun *et al.,* 2013). Finally, the processed SLAF tags were sequenced using an Illumina HiseqTM 2500 (Illumina, Inc; San Diego, CA, USA).

### Evaluation of data quality

The raw output produced by the Illumina HiseqTM 2500 was further analyzed for each sample using the software “Dual-index” (Kozich *et al.,* 2013). After removing adapter sequences, raw short reads were assessed by calculating GC contents and Q30 (Q = -10 × log_10_^e^; indicating a 0.1% chance of an error). All SLAF reads were clustered based on sequence alignments using the BLAT software (Kent, 2002). Polymorphic SLAF tags were determined by comparisons of sequence variation between different oil palm individuals. SLAF tag sequences were mapped to the whole genome of *Elaeis guineensis* using the Burrows-Wheeler alignment tool (BWA) software (Li and Durbin, 2009).

### Identification of SNP markers

SNP markers were identified for polymorphic SLAF tags using two software programs: GATK (McKenna *et al.,* 2010) and SAMtools (Li *et al.,* 2009). SNP markers identified consistently between the two methods were considered to be reliable. Finally, SNPs matching the criteria of a minor allele frequency (MAF) of more than 0.05 and integrity above 1 were selected for subsequent analysis.

#### Identification of genes, putative SSRs and retrotransposons

The gene annotation result were downloaded from the National Center for Biotechnology information in *Elaeis guineensi*s. The mining of retrotransposons (TEs) in Elaeis guineensis was done using RepeatMasker (Smit et al., 2013/2015). The SSR analysis software Msatfinder (https://github.com/knirirr/Msatfinder) was used to identify all possible mono-, di-, tri-, tetra-, penta-, and hexa-nucleotide SSRs with a minimum set of 12, 4, 4, 4, 4, and 4 repeats, respectively (Thurston and Field, 2005)

### Extraction and measurement of fatty acid contents in oil palm mesocarp tissues

Three fruits per oil palm individual (three biological replications) were harvested, and fatty acid extraction and analyses for each mesocarp tissue was performed in triplicate (three different extractions as technical replicates). Approximately 5 mg of mesocarp was used for extracting fatty acid according to methods described in Li-Beisson *et al.* (2013). Subsequently, fatty acid composition was examined and measured using gas chromatography (Agilent DB-23, 30 m×250). The heating procedure was carried out as follows: the initial temperature was set as 180 °C, followed by temperature increases up to 220 °C at a frequency of 10 °C per minute. The gas pressure used was 17.392 psi. The nine values obtained per oil palm individual were averaged for subsequent association mapping.

### Population structure and linkage disequilibrium analysis

Bayesian clustering was applied to analyze the population structure of 200 oil palm individuals using the software STRUCTURE (Pritchard *et al.,* 2000). Based on the same set of SNPs, the number of subgroups (*K*) was predicted from 1 to 10, and the number of ancestors was determined according to the position of the minimum value, with error rate obtained from 5-fold cross-validation. Maximum likelihood estimates for the ancestry proportion from each K subgroup for each accession were calculated.

Linkage disequilibrium (LD) across the genome of *Elaeis guineensis* was calculated using the software package TASSEL 4.0 (Bradbury *et al.,* 2007). LD decay for each chromosome was evaluated at a cut-off value of *r^2^* = 0.1. The r^2^ value for a marker distance of 0 Kb was assumed to be 1.

### Linkage disequilibrium blocks

The LD blocks present in the 200 oil palm individuals were estimated using the HAPLOVIEW v4.2 software. The number and size of the LD blocks present on every chromosome were calculated according to established methods (Barrett *et al.,* 2005).

### Association mapping

In order to reduce spurious associations between alleles and phenotypic variation, mixed linear models (MLM) were used. Fixed effects were computed using a Q (population) value matrix, and random effects were computed using a K (Kinship) matrix. The Q+K value matrix was added into the MLM model. The Q matrix was obtained using the STRUCTURE software (Pritchard *et al.,* 2000), and the K matrix (genetic relationship between the 200 oil palm individuals) was obtained using the software SPAGeDi (Hardy *et al.,* 2002). P values between SNP markers and fatty acid content were computed using the following formula:

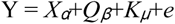

where Y represents the phenotype value, X represents the genotype, *Q_β_* refers to fixed effects and *K_μ_* refers to random effects. Quantile-Quantile plots (Q-Q plots) were drawn using the “GGplot2” software R package (Ginestet, 2011), and the Manhattan plot was drawn using the “qqman” software package (Turner *et al.,* 2011).

### Fatty acid content candidate gene prediction

All genes in LD blocks (*r^2^* > 0.6) containing SNPs significantly associated with traits were identified for further candidate gene selection (Raman *et al.,* 2015). If SNP markers significantly associated with traits were located outside LD blocks, genes in the 200 Kb flanking regions of the linked markers were selected for further analysis. Amino sequences of these selected genes were used as queries to blast against the *Arabidopsis* protein database in order to predict the potential function of candidate genes. All candidate genes were selected based on gene ontology (GO) terms related to fatty acid biosynthesis and metabolism and all transcription factors were selected based on Clusters of Orthologous Group of proteins (COG) within SNP-tagged genome regions.

### Transcriptome data downloaded and RPKM calculation

We downloaded transcriptomic raw read data from the SRA (Short Read Archive) database, including SRR851069 (mesocarp 10 weeks after anthesis), SRR851067 (mesocarp 15 weeks after anthesis), SRR190699 (mesocarp 17 weeks after anthesis), SRR190700 (mesocarp 19 weeks after anthesis), SRR190701 (mesocarp 21 weeks after anthesis), SRR190702 (mesocarp 23 weeks after anthesis), SRR851070 (kernel 10 weeks after anthesis), SRR851068 (kernel 15 weeks after anthesis), SRR190703 (leaf), SRR851071 (root), SRR851103 (shoot), SRR851101 (female flower), and SRR851099 (pollen). RPKM (reads per kb per million reads) were used to calculate gene expression levels using the following formula: (Mortazavi *et al.,* 2008)

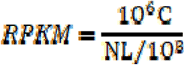

where C is the number of reads that aligned exclusively with one expressed sequence, N is the total number of reads that aligned with all expressed sequences, and L is the number of bases in the corresponding coding sequence.

### Vector construction and transformation for over-expression of *EgFatB*

Primers used for *EgFatB* gene cloning were designed using the Snapgene Viewer software (Table S4). PCR amplifications were performed in 50 µl reaction mixtures containing a 500 ng cDNA sample from the oil palm mesocarp, 1 × PCR buffer, 2 mM MgCl_2_, 5 U TaqDNA polymerase (TaKaRa, China), 0.5 μM of each primer and 0.2 mM dNTP mix, with the following program: denaturation for 30 seconds at 98°C, 30 cycles of 98°C for 10 seconds, 30 seconds at 55°C and 1 minute at 72°C for elongation, with a final extension of 5 minutes at 72°C. Subsequently, PCR products were electrophoretically visualized on 1% agarose gel and recombined into the *pBinGlyRed3* vector (containing DsRed as a reporter gene and 35S as a promoter), which was then digested with EcoR I and XhoI. The constructed vectors were transformed into competent *E. coli* cells (line Dh5l) and inserts validated by sequencing. A plasmid with the correct sequence insert was transformed to *Agrobacterium GV3101* and then transformed into *Arabidopsis thaliana* using an *Agrobacterium* mediated *in planta* transformation approach (Clough *et al.,* 1998). Positive transgenic *Arabidopsis* were confirmed by detection of red autofluorescence in seeds and by PCR validation of targeted genes. The control and transgenic Arabidopsis were grown under 16 hours light/8 hours dark photoperiod at 25 °C.

## Results

### SLAF-seq of 200 oil palm individuals

Reads derived from SLAF sequencing of the two hundred oil palm individuals were filtered and adaptor sequences removed, resulting in 908.37 Mb of reads with a Q30 of 90.15% and a GC content of 40.79%. A total of 357 378 SLAF tags were obtained from the 200 oil palm individuals, with an average coverage of 10.23 × per sample. SLAF tags were mapped to the reference oil palm genome using the software “BWA”, and 249 457 SLAF tags containing polymorphic SNPs were detected among the 200 oil palm individuals (Table S1 and Figure 1). Of these SLAF tags containing polymorphic SNPs, 43.62% (108 820) were located on the 16 assembled chromosomes of the oil palm reference genome, similar to the percentage 42.28% of the total assembled genome sequence of oil palm present in the chromosome assemblies (642.5 Mb out of 1.535 Gb). The number of SLAF tags per chromosome varied from 5 673 (Chr16) to 17 521 (Chr1). A total of 5 870 684 SNPs were identified among the 200 individuals of oil palm, of which 1 261 501 SNPs were identified by both GATK and SAMtools with MAF > 0.05 and integrity > 1, and were subsequently used as SNP markers. The raw datas of 200 oil palm individuals’ SLAF-seq had been deposited onto European Nucleotide Archive (https://www.ebi.ac.uk/ena). The bioproject numer is PRJEB26466.

**Figure 1.**
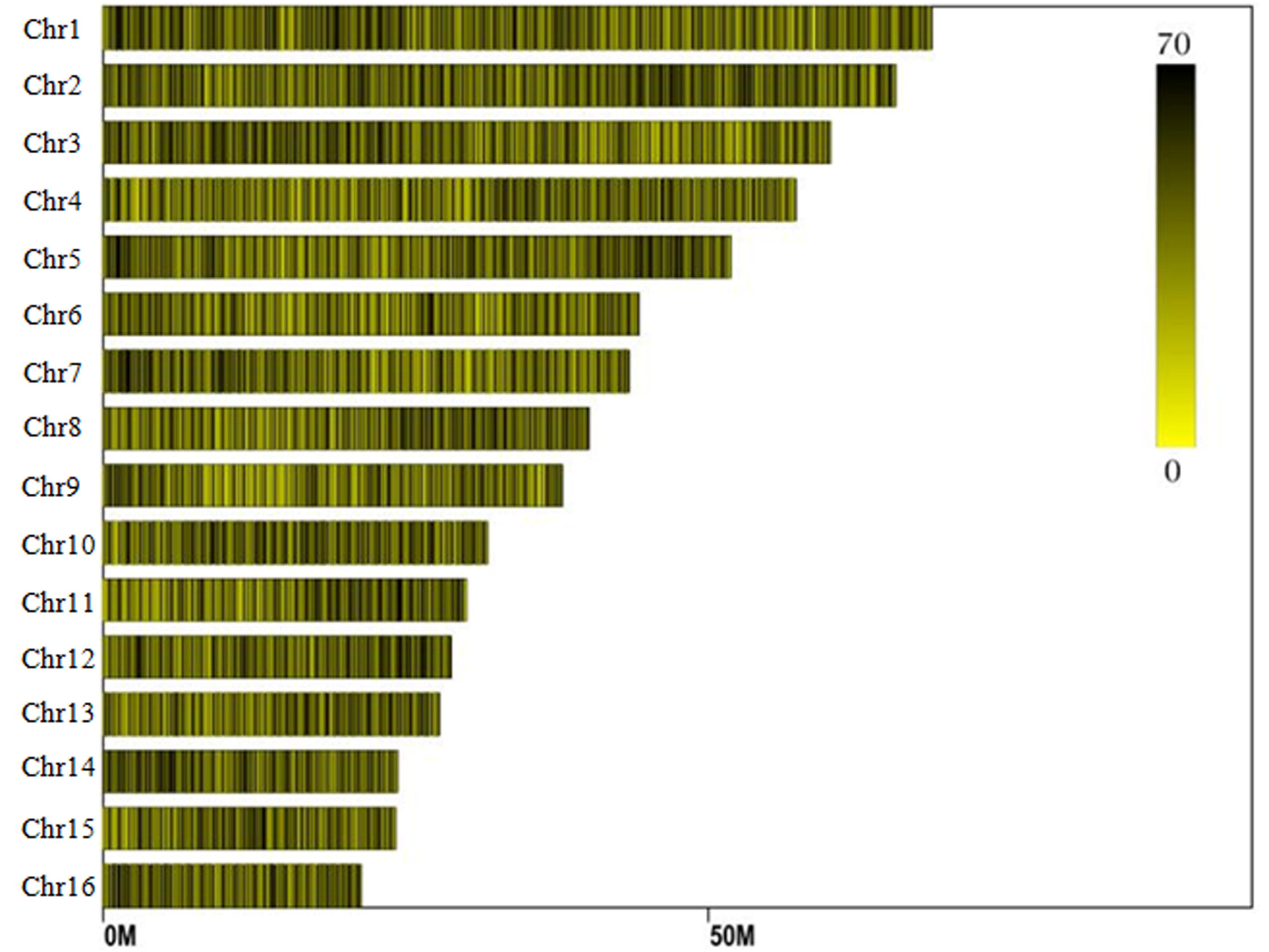
Distribution of SLAF (Specific Locus Amplified Fragment) tags across the chromosomes of the oil palm reference genome.

### Genomic distribution of the SNPs in *Elaeis guineensis*

From the oil palm genome annotation results, 22 957 protein coding genes are distributed on the 16 assembled chromosomes, with gene number per chromosome ranging from 2662 (Chr1) to 812 (Chr16) (Date from NCBI). Of the 1.2 million SNP markers located in the 249 457 polymorphic SLAF tags, 17.81% were located within genic regions of 5064 genes (7.38 SNP markers per gene on average). Among these, the largest number of SNP markers (11 319) were located in 2662 genic regions on the chromosome 1, with an average of 4.25 SNP markers per gene, followed by chromosome 2 (10 960 SNP markers in 1769 genes) and chromosome 3 (8 491 SNP markers in 1513 genes).

For 1,261,501 SNP markers identified in 200 oil palm individuals, 40.8% of SNPs were evenly distributed across the 16 chromosomes (Figure 1). The number of SNP markers per chromosome varied from 17 554 SNPs (Chr16) to 52 023 SNPs (Chr11), with an average of 32 168 SNP markers per chromosome.

Analysis of the genomic distributions of SNP markers, microsatellites, transposable elements (TEs) and predicted genes revealed a negative correlation between the distribution of SNP markers and genes (correlation coefficient: -0.1355) (Figure 2). However, a positive correlation was detected between the distribution of SNP markers and TEs (correlation coefficient: 0.3443).

**Figure 2.**
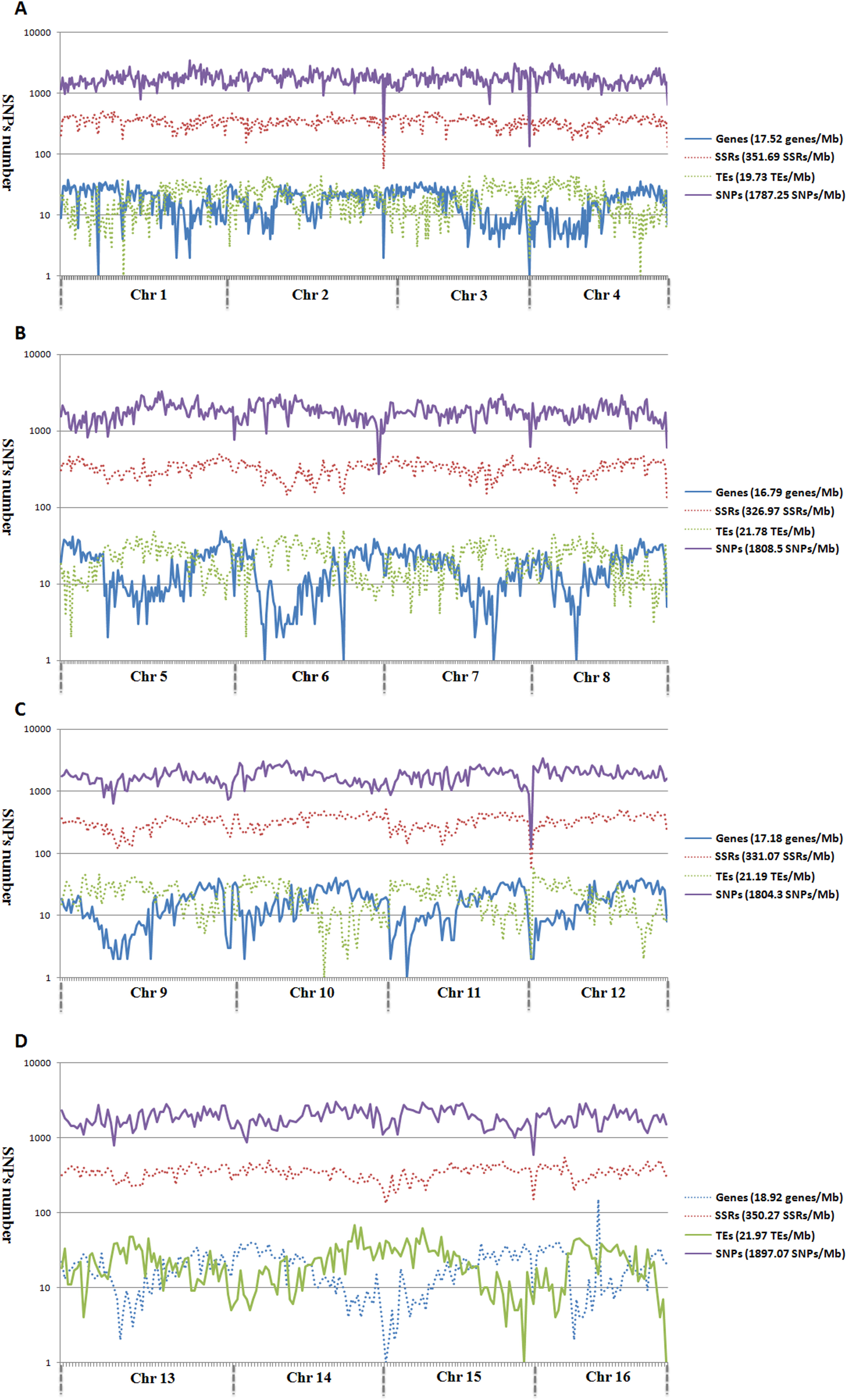
Genomic distribution of genes, microsatellites, TEs, and SNPs across the 16 assembled chromosomes of oil palm

In order to ascertain the relationship between SNP and retrotransposon (TEs) distributions, the number of SNP markers in the 1 Kb to 10 Kb flanking regions of TEs was calculated (Figure S1). SNP distances to TEs were positively correlated with the SNP number for all chromosomes except for Chr 1 and Chr 5 (Figure S1). Meanwhile, the total number of SNP markers gradually decreased with increasing distance from TEs.

### Genetic structure of the experimental population of oil palm

Population structure analysis based on the SNP markers divided the 200 oil palm individuals into five subgroups based on cross validation (CV) errors (Figure 3).The analysis results showed that different populations cannot be divided by geographical origin. In particular, the individuals from Southern China and Costa Rica were distributed across the five subpopulations. However, the second population contained most (71%, 57 out of 80) of the individuals from Malaysia, while, the fifth population contained nearly half (49%, 44 out of 90) of the individuals from Malaysia.from the plateau region of Costa Rica. Most (5, 62.5%) of the remaining individuals were assigned into the fourth population. The firist population were totally mixed by geography, including 2 individuals from Southern China, 10 from Malaysia, 12 from the plateau region of Costa Rica, and 3 from Africa.

**Figure 3.**
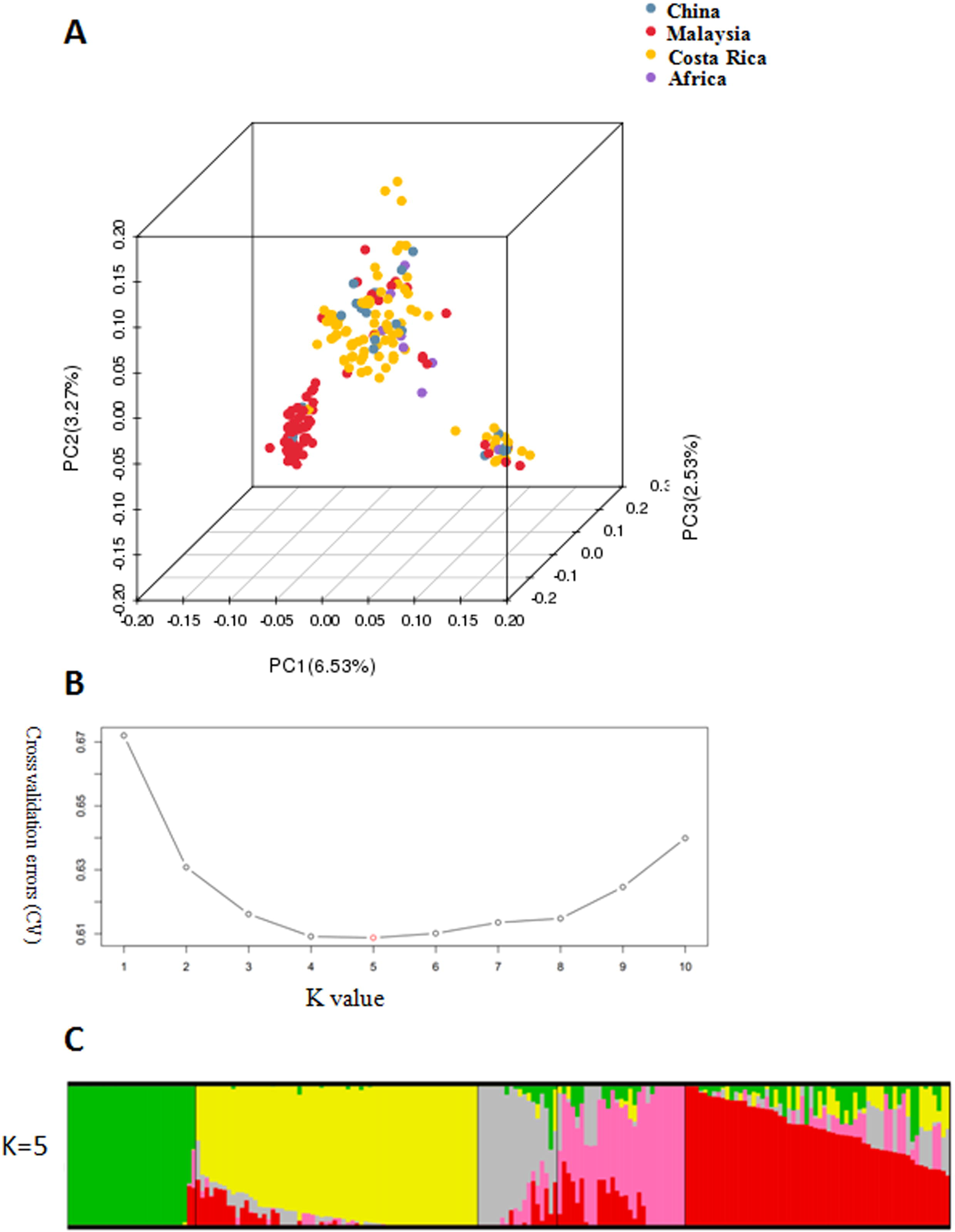
Population structure of 200 oil palm individuals from four different geographical locations. (A) Principal component analysis (PCA) of 200 oil palm individuals, showing the total variance explained by the first (PC1: 6.53%), second (PC2: 3.27%) and third (PC3: 2.53%) principal components. (B) Cross validation (CV) errors suggest that the 200 oil palm individuals were divided into five true genetic populations. (C) Population structure of the 200 oil palm individuals.

Genetic diversity of the 200 oil palm individuals was analyzed (Table S2). The average observed heterozygous heterozygosity and polymorphism information content is was 0.243 and 0.274, respectively. The group of population comprised of 80 oil palm individuals from Malaysia showed the highest observed allele number (1.996) and observed heterozygous heterozygosity (0.25), which havebut had a similar diversity level of genetic diversity with the population group of 90 *Elaeis guineensis* individuals from Costa Rica. Relatively low diversity levels were detected in oil palm individuals from Africa and China.

In order to evaluate extent and pattern of LD among the 200 oil palm individuals, the LD decay was analyzed, with an *r*^2^ threshold of 0.1. The average physical distance for LD decay was 14.5 Kb genome-wide (Figure S2). However, different LD decay distances were observed for different chromosomes, ranging from 3.3 Kb (Chr15) to 20.0 Kb (Chr5).

### Haplotype blocks based on linkage disequilibrium

Haplotype blocks were evaluated based on LD estimation across the 200 oil palm individuals. A total of 9 286 conserved haplotype blocks were detected among the 200 oil palm individuals, spanning 233.77 Mb (37.11% of the assembled reference genome). Among haplotype blocks, 54.77% varied in size from 0 to 1 Kb, while 6.98% were more than 100 Kb in size. The haplotype block number varied from 280 (Chr16) to 1 048 (Chr2), with an average of 580 haplotype blocks per chromosome (Table S3). Haplotype block size per chromosome varied from 7.79 Mb (chromosome 16) to 23.69 Mb (chromosome 1), with an average of 13.98 Mb per chromosome (Table S3). The percentage of each chromosome containing haplotype block size varied from 30% in chromosome 14 to 36.6% in chromosome 15, with an average haplotype block percentage of 33.79% per chromosome.

### Fatty acid composition in the experimental oil palm population

In order to perform association analysis between SNP markers and fatty acid compositions, we investigated the fatty acid composition of each the 200 oil palm individuals from different geographical origins (Figure 4). Among the 200 oil palm individuals, palmic acid content varied from 31.3% to 48.8% with an average of 42.0%, oleic acid content varied from 31.3% to 50.1%, linoleic acid content varied from 7.1% to 18.5%, and total oil content varied from 29.8% to 70.3%.Palmic acid content (C16:0) was significantly negatively associated with stearic acid (correlation coefficient: -0.627; P < 0.001) and oleic acid (correlation coefficient: -0.657; P < 0.001). Oleic acid content was also significantly negatively correlated with linoleic acid content (correlation coefficient: -0.689: P < 0.001).

**Figure 4.**
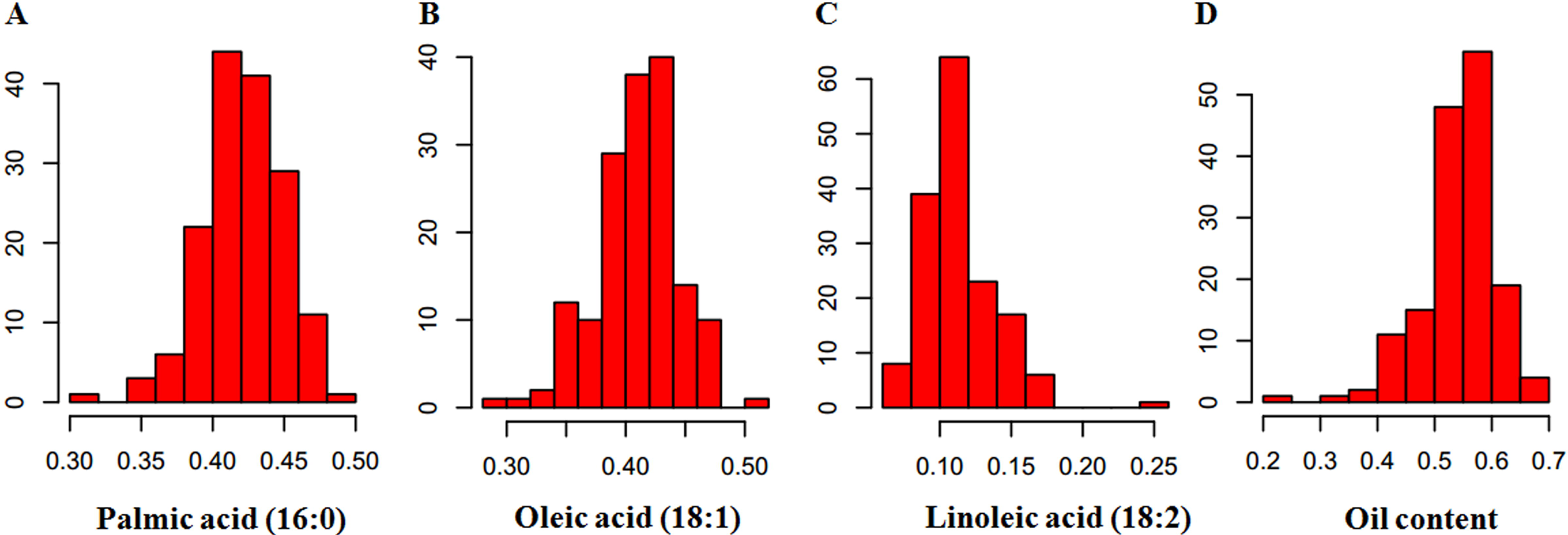
Proportion of fatty acid content in the mesocarp of 200 oil palm individuals: (A) palmic acid content (16:0); (B) oleic acid content (18:1); (C) linoleic acid content (18:2) and (D) oil content.

### Genome-wide associations between SNPs and fatty acid content

In order to detect associations between allele variation and fatty acid content, mixed linear models (MLM) were used for GWAS to control for the presence of population structure in the analyzed oil palm population, and SNPs significantly associated with traits were displayed on Manhattan plots (Figure 5 and Figure S3). The MLM analysis revealed a total of 274 SNP markers (P < 1e^-4^) significantly associated with fatty acid content, including palmic acid content (85 SNPs), oleic acid content (35 SNPs), linoleic acid content (130 SNPs), and total oil content (98 SNPs) (Figure 7). These SNP markers were distributed across all oil palm chromosomes except Chr11. The largest number of associated SNP markers (27) was located on Chr1, followed by Chr5 (21 SNP markers) and Chr4 (15 SNP markers). Thirteen SNP markers were associated simultaneously with total oil content, oleic acid content (18:1) and linoleic acid content (18:2). A further forty six SNP markers were significantly associated with both oil content and linoleic acid content (18:2), and thirteen SNP markers were significantly associated with both oil content and oleic acid content (18:1).

**Figure 5.**
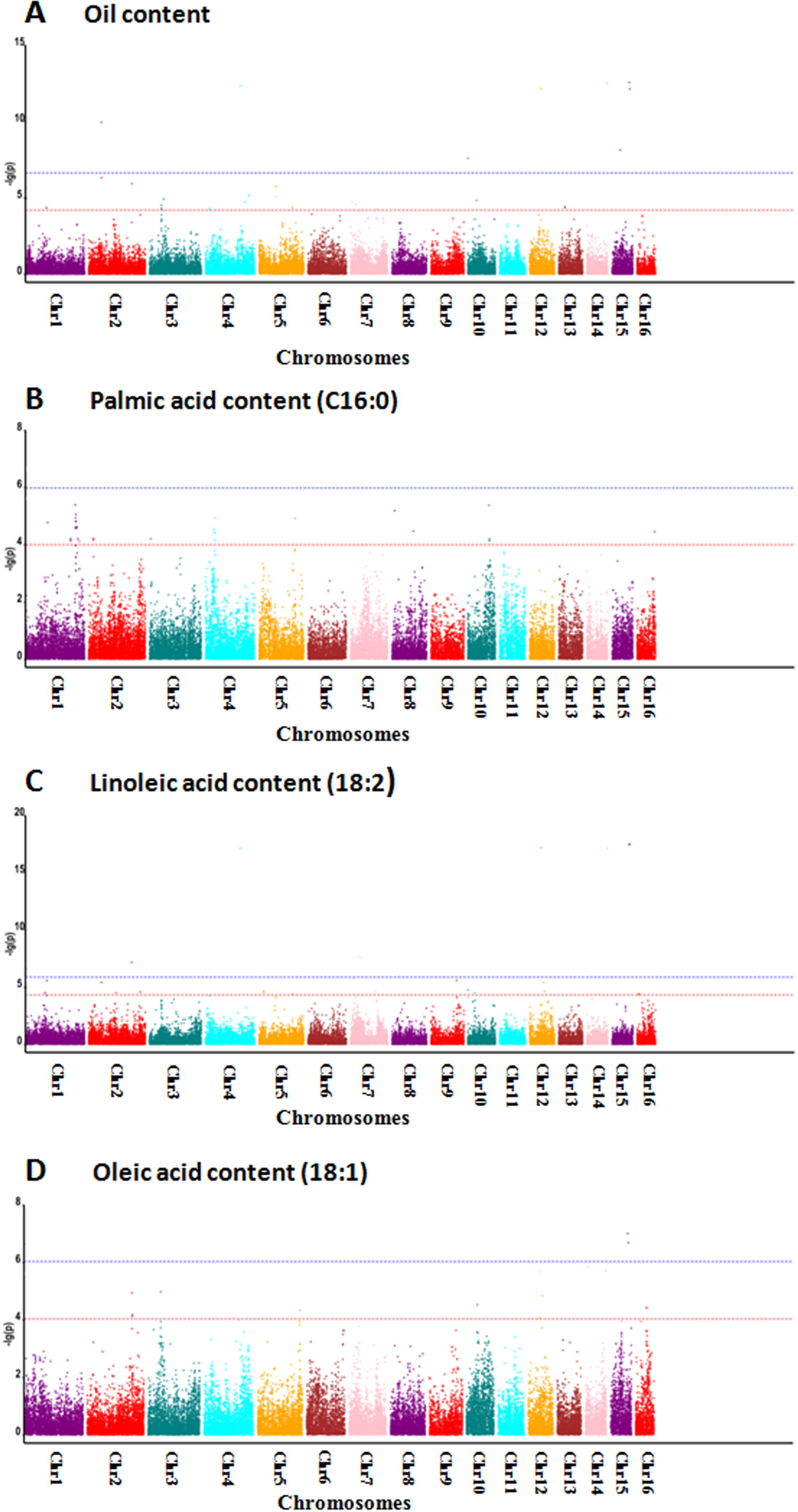
Genome-wide associations between SNP markers and fatty acid contents in oil palm (*Elaeis guineensis*) using mixed linear models: (A) Manhattan for oil content; (B) Manhattan for palmic acid content (C16:0); (C) Manhattan for linoleic acid content (18:2); and (D) Manhattan for oleic acid content (18:1).

### Identification of candidate genes related to fatty acid composition in *Elaeis guineensis*

Of the 274 SNP markers significantly associated with fatty acid content, 69 were located in different LD blocks (r^2^ > 0.6), while the other SNP markers were not located in defined LD blocks. A total of 1 148 candidate genes were identified in the flanking regions of 129 SNP markers significantly associated with different fatty acid contents. Based on GO annotation results, 44 candidate genes were involved in the fatty acid biosynthesis, metabolism and the tricarboxylic acid cycle pathway. These included two acyl-ACP thioesterase B genes (FatB, involved in the termination of fatty acid chain), stearoyl-ACP desaturase (SAD, involved in fatty acid dehydrogenation), oleate desaturase (FAD, involved in fatty acid dehydrogenation), ketoacyl-CoA reductase (KCR, involved in fatty acid elongation), ketoacyl-CoA synthase (KCSII, involved in fatty acid elongation), ketoacyl-ACP synthase (KAS, involved in fatty acid elongation), AP2/EREBP transcription factors (WRI1, involved in fatty acid synthesis), lipid transfer protein (LTP), Acyl-CoA: diacylglycerol acyltransferase (DGAT), malonyl-CoA synthase (MCS), among others (Supplementary Table 1).

### Candidate gene expression in different oil palm tissues

The expression of candidate fatty acid biosynthesis and metabolism genes in different tissues and in different mesocarp developmental stages was calculated using RPKM values based on the method of Mortaavi et al. (2008). Of the 44 candidate gene involved in fatty acid biosynthesis and metabolism identified based on GO annotation results, 33 (75%) showed comparatively high expression levels in the mesocarp tissues (Figure 6 and Table S4).

**Figure 6.**
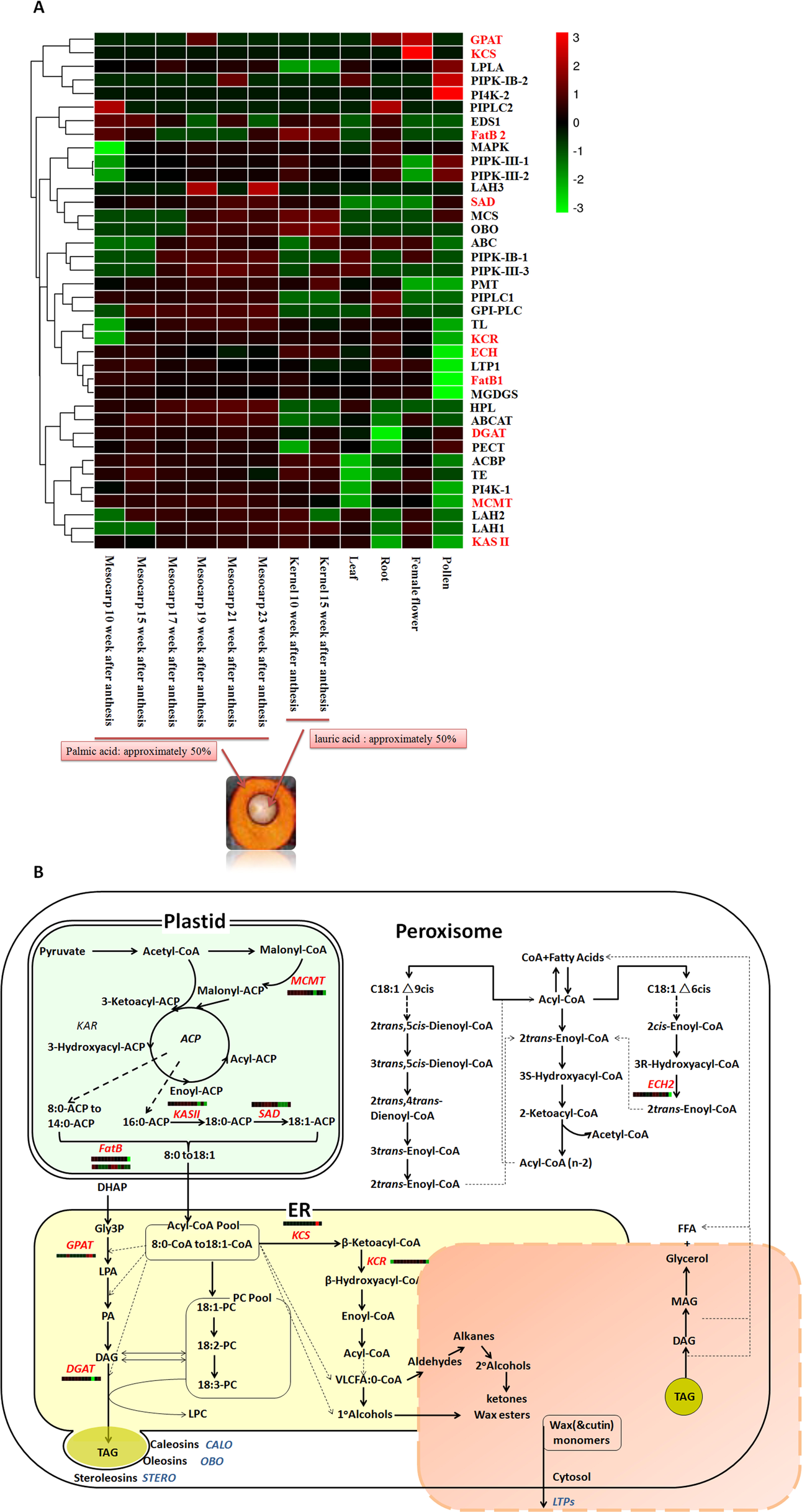
Expression patterns of candidate genes adjacent to SNP markers significantly associated with fatty acid biosynthesis and metabolism in different oil palm tissues. (A) Heatmap of expression of the 44 candidate genes in different tissues of oil palm. (B) The fatty acid biosynthesis and metabolism pathway, with candidate genes identified in the pathway marked in red.

Example genes of particular interest included MCMT (ACP-malonyl transferase), KASII (beta-ketoacyl-acyl carrier protein synthase II), SAD (stearoyl-ACP desaturase) and DGAT (acyl-CoA:diacylglycerol acyltransferase). The MCMT gene plays an important role in fatty acid biosynthesis and can transfer malonyl from malonyl-CoA to an acyl carrier protein and release free CoA. The MCMT gene also had a high expression level in the late developmental stages of the mesocarp, with RPKM values of 6.52 at 10 weeks post-anthesis, 102.13 at 21 weeks post-anthesis, and 154.19 23 weeks post-anthesis (Figure 8a and Table S?). Meanwhile, this gene was also highly expressed in the kernel (RPKM values 62.2 and 24.52 at 10 and 15 weeks after anthesis), but lowly expressed in other tissues (RPKM value: 4.05, 0, 3.71, 4.94, and 0 in leaf, root, shoot, female flower, and pollen respectively). KASII (beta-ketoacyl-acyl carrier protein synthase II) is involved in the elongation of palmitoyl-ACP (C16:0-ACP) to stearol-ACP (C18:0-ACP). This gene had comparatively high expression levels in different developmental stages of the mesocarp and kernel. However, this gene also had high expression levels in leaves and female flowers. SAD (stearoyl-ACP desaturase) can catalyze desaturation of stearoyl-ACP to form oleoyl-ACP. This SAD gene had a high expression level at 21 (RPKM value: 66.1) and 23 (61.48) weeks post-anthesis in the mesocarp, but low expression levels in other tissues. Candidate DGAT (acyl-CoA:diacylglycerol acyltransferase) involved in TAG synthesis also had a higher expression level in the six developmental stages of the mesocarp compared to in other tissue types.

Two FatBs (Acyl-ACP thioesterase B) were detected close to one significant SNP (Eg-chr6-43621940) and two significant SNPs (Eg-chr7-1731788 and Eg-chr7-1731870) respectively. These two FatB genes are hereafter referred to as *EgFatB1* and *EgFatB2*. FatB genes have been reported to be involved in termination of saturated fatty acyl-ACP. Interestingly, *EgFatB1* had higher expression levels in different developmental stages of the mesocarp, especially at 10 weeks and 15 weeks post-anthesis. However, *EgFatB2* only had high expression levels at 10 weeks (RPKM value: 187.61) and 15 weeks (92.44) post-anthesis in the kernel.

As well as the 44 candidate genes involved in fatty acid biosynthesis and metabolism, relative expression patterns of the remaining 1104 candidate genes located in the same haplotype block or within 200 Kb of signifcantly associated SNP markers were also analyzed. Among these candidate genes, 136 were highly expressed in different developmental stages of the mesocarp and kernel, and 10 candidate genes showed high expression only in the kernel. Meanwhile, 21 candidate genes had very high expression levels (RPKM > 100) in either the mesocarp or kernel. Of these 21 candidates, six were annotated as uncharacterized proteins, while five were annotated as involved in plant hormone response and regulation, including two shaggy-related kinase orthologues involved in the brassinosteroid signal transduction pathway, two IAA1 protein orthologues and one gibberellin–beta-dioxygenase orthologue. The remaining 10 candidates were annotated as transcription factors (3), amino acid phosphorylators (2), and as TIP2-1 involved in water transport (2).

### Functional validation of *EgFatB1* and *EgFatB2*

*EgFatB1* and *EgFatB2* were cloned and linked to pBinGlyRed3 vectors which contained a red autofluorescence gene (DsRed3) as a reporter gene and a 35S promoter. The *EgFatB1* and *EgFatB2* over-expression vectors were then transformed into *Arabidopsis thaliana*. Positive transformants were screened in the T1 generation by detection of red autofluorescence in the seeds and by PCR validation. T2 generations of positive transgenic plants were investigated for fatty acid content. Compared to wild-type *Arabidopsis* seeds, the palmic acid content in *EgFatB1* transgenic plants increased from 7% to 32%. Stearic acid content also increased from 3% to 6%, oleic acid content decreased from 21% to 10%, and linoleic acid decreased from 32% to 21% (Figure 7). Overall, over-expression of *EgFatB1* significantly increased saturated fatty acid content in *Arabidopsis* seeds.

**Figure 7.**
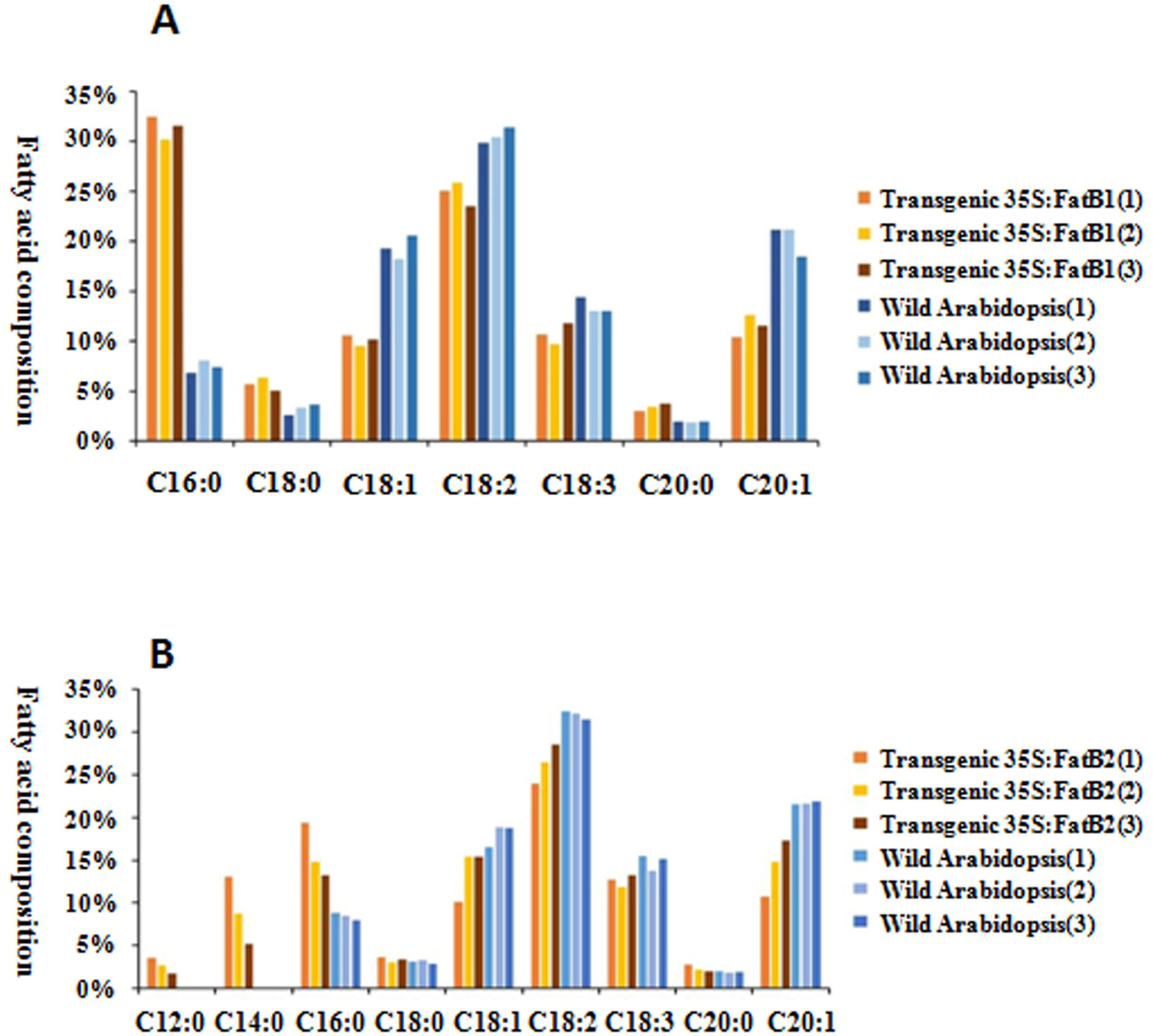
Fatty acid composition in transgenic 35S:*EgFatB1*, transgenic 35S:*EgFatB2* and wild-type *Arabidopsis thaliana*. (A) Comparative analysis of fatty acid content between transgenic 35S:*EgFatB1* and wild-type *Arabidopsis*; (B) Comparative analysis of fatty acid content between transgenic 35S:*EgFatB2* and wild-type *Arabidopsis*

The over-expression of *EgFatB2* in *Arabidopsis thaliana* also significantly increased palmic acid content from 10% to 20%. Oleic acid content was also increased from 11% to 18%, and linoleic acid decreased from 16% to 12%. Although wild-type *Arabidopsis* plants do not contain lauric or myristic acid, these fatty acids were detected in the 35S:*EgFatB2* transgenic *Arabidopsis* (Figure 7)

## Discussion

Oil palm (*Elaeis guineensis*) is a crop of major importance for human nutrition, with tens of millions of tonnes of palm oil consumed every year. However, oil palm breeding, which could potentially improve the nutritional value of palm oil for human consumption (e.g. by lowering saturated fat content) is hindered by the lack of genetic and genomic resources available for this crop. Using SLAF-seq technology, we genotyped a diversity panel of 200 oil palm individuals from four countries to provide 1.2 million genome-wide SNPs, and associated these SNPs with fatty acid content to identify dozens of candidate genes involved in fatty acid biosynthesis and metabolism pathways. Two genes with tissue-specific expression profiles suggestive of their role in fatty acid composition changes (*EgFat1B* and *EgFat2B*) were validated using *Arabidopsis* transformation: over-expression of both genes resulted in increased saturated fat content in *Arabidopsis* seeds. Interestingly, over-expression of *EgFat2* also led to the production of lauric and myristic acid in *Arabidopsis* seeds, despite the fact that these fatty acids do not naturally occur in *Arabidopsis*. These genes provide an excellent target for future gene knock-out or mutation breeding approaches to recover oil palm mutants with lower saturated fat content. Our results provide a comprehensive set of genetic resources for oil palm breeding, particularly in improving the fatty acid composition of this important oil crop for human nutrition.

Single nucleotide polymorphisms have a much higher density in the genome than other molecular markers, and as such are highly suitable for use as markers in population structure and linkage disequilibrium analysis. With the rapid development and application of next-generation sequencing (NGS) technology, various reduced-representation genome sequencing approaches have been developed and applied in various species for SNP genotyping. These include genotype-by-genotype (GBS) (Elshire *et al.,* 2011), RAD (restriction-site associated DNA sequencing, Bus *et al.,* 2012), RRL (reduced representation libraries, Tassell *et al.,* 2008) and SLAF-seq, which is an accurate and cost-effective sequence-based SNP identification method (Sun *et al.,* 2013; Zhang *et al.,* 2013). Chen *et al.* (2014) reported the first use of SLAF-seq technology, developing 89 specific and stable molecular markers in *Thinopyrum elongaum*. Zhang *et al.* (2016) reported construction of a high-density genetic map harboring 5521 single nucleotide polymorphism markers (SNPs) and its application to quantitative trait loci (QTL) analysis for boll weight in upland cotton (*Gossypium hirsutum*). Zeng *et al.* (2017) reported a genome-wide association study using 2 309 777 SNP markers based on SLAF-seq technology and their association with resistance to rust disease in orchardgrass. Geng *et al.* (2016) used SLAF-seq to develop 1933 SNP markers in *Brassica napus*, subsequently identifying four SNP markers significantly associated with seed weight. Hence, SLAF-seq technology has been validated to be an efficient method for SNP marker identification, and one which has been widely applied in various crop species. In our study, a total of 1 261 501 SNP markers were identified using SLAF-seq, with an average SNP density of 70 SNPs/100 Kb, and an average physical distance between adjacent SNP markers of 0.5 Kb. As we ascertained the average LD decay distance to be 14.51 kB in oil palm, the high density and numbers of SNPs obtained via SLAF-seq in our study allowed precise haplotype map construction and high-resolution association analysis.

Unexpectedly, our results revealed little correlation between geographic origin and genetic relationship in our oil palm diversity set, although five genetic subpopulations were identified. As well, oil palm individuals from Malaysia and Costa Rica showed comparatively high genetic diversity when compared to oil palm individuals from Africa and China. This was surprising for the African individuals, as oil palm is thought to have originated along the coast of the Gulf of Guinea in Western and Central Africa, but may reflect sampling bias due to the lower number of samples obtained from Africa. The expansion of oil palm as an industrial and food crop began in Africa and Southern Asia (especially Malaysia) during the colonial period. This tropical species was initially introduced into Malaysia as an ornamental tree in the 1870s, but by the 1920s was widely cultivated for oil production: Malaysia currently produces 40% of all palm oil world-wide. In the 1960s the oil palm variety “Dura” was introduced from Malaysia into southern China (Hainan province), but initial plantation trials failed due to poor adaptability of the selected variety to the climate and soil environment in southern China. Later, efforts were made to collect oil palm germplasm from different tropical regions and test their adaptability to the climate and soil conditions in southern China, as well as to evaluate their agronomic traits (Cao *et al.,* 2012). However, our results suggest that oil palm in southern China still has relatively low genetic diversity, suggesting most oil palm resources are probably derived from the same “Dura” variety introduced from Malaysia in the 1960s. Overall, very little is known about the genetic population structure of oil palm, which hinders germplasm collection and pre-breeding efforts: our results provide a useful resource for future research in this direction.

Marker-assisted selection has major potential to speed up breeding of oil palm cultivars with improved fatty acid composition (e.g. decreased palmic acid content) in order to increase the market and nutritional value of palm oil. Identifying trait-associated markers is a prerequisite for marker-assisted selection. In previous research, QTL mapping has been used to identify markers linked to agronomic traits based on biparental cross populations of oil palm. The first genetic linage map was constructed using restriction fragment length polymorphism (RFLP) markers (Rance *et al.,* 2001). QTL mapping was subsequently used to decipher the genetic bases of yield components (Rance *et al.,* 2001; Billotte *et al.,* 2010), fatty acid composition (Singh *et al.,* 2009; Montoya *et al.,* 2013), and sex ratio (Ukoskit *et al.,* 2014), among other traits. These QTL mapping experiments involved populations produced from one to four parents. Montoya *et al.* (2013) reported identification of nineteen QTLs associated with fatty acid composition in palm oil. However, the confidence interval of these QTLs ranged from 6 cM to 30 cM, or approximately 6 – 30 Mbp of physical genome sequence in oil palm. Therefore, utilization of these markers for marker-assisted selection is not possible due to the large physical and genetic distance between the linked markers and underlying target genes. Data produced in previous mapping studies suffers from two major shortcomings in terms of suitability for marker-assisted selection: (1) low-density molecular markers were used in the construction of the linkage map; and (2) relatively low recombination frequencies in biparental cross populations were obtained compared to in natural populations. In our study, genome-wide association analysis revealed a total of 274 SNP markers significantly associated (P < 1e^-4^) with four fatty acid traits, distributed across 15 of the 16 oil palm chromosomes. Although most previous studies are not directly comparable to our results in terms of gene identification, Montoya *et al.* (2012) reported 19 QTL for fatty acid composition using a backcross population of *Elaeis oleifera* with *Elaeis guineensis*: underlying these QTL were acyl-ACP thioesterase A and Stearoyl-ACP desaturase genes, which were also found as gene candidates adjacent to SNP markers significantly associated with fatty acid in our study.

## Acknowledgements

This work was supported by the Scientific and Technological Cooperation Projects of Hainan province (No. KJHZ2015-06) and DFG Emmy Noether grant MA6473/1-1.

## Supplementary Information

Figure S1 The number and distribution of SNP markers in the 1Kb to 10Kb flanking region of TEs in chromosomes 1–6 (A), chromosomes 7-12 (B), chromosomes 13-16 (C) and averaged across all chromosomes (D). Horizontal coordinates represent the flanking regions of TEs. Vertical coordinates represent the number of SNP markers.

Figure S2 Linkage disequilibrium (LD) decay of the oil palm genome and across different chromosomes

Figure S3 Genome-wide associations between SNP markers and fatty acid contents in oil palm (*Elaeis guineensis*) using mixed linear models: (A) Quantile-quantile plots for oil content; (B) Quantile-quantile plots for palmic acid content (C16:0); (C) Quantile-quantile plots for linoleic acid content (18:2); and (D) Quantile-quantile plots for oleic acid content (18:1).

Table S1 The distribution of SLAF (Specific Locus Amplified Fragment sequencing) tags produced across the oil palm (Elaeis guineensis) genome

Table S2 TGenetic diversity analysis of different populations of 200 oil palm individuals from different geographic origins, including showing observed number of allele numbers, expected number of allele snumber, observed number of heterozygous numberloci, expected number of heterozygous numberloci, and polymorphism information content.

Table S3 Distribution of haplotype blocks in the genome of *Elaeis guineensis*

Table S4 Primers’ sequences used to amplify the full CDS sequence of FatB1 and FatB2

## References

Atwell S, Huang YS, Vilhjalmsson BJ, et al. 2010. Genome-wide association study of 107 phenotypes in a common set of *Arabidopsis thaliana* inbred lines. Nature 465, 627–631.

Barrett JC, Fry B, Maller J, Daly MJ. 2005. Haploview: analysis and visualization of LD and haplotype maps. Bioinformatics 21, 263–265.

Billotte N, Jourjon MF, Marseillac N, Berger A, Flori A, Asmady H, et al. 2010. QTL detection by multi-parent linkage mapping in oil palm (*Elaeis guineensis* Jacq.). Theoretical and Applied Genetics 120, 1673–1687.

Bonaventure G, Bao X, Ohlrogge J, Pollard M. 2004. Metabolic response to the reduction in palmitate caused by disruption of the FatB gene in Arabidopsis. Plant Physiology 135, 1269.

Bradbury PJ, Zhang Z, Kroon DE, Casstevens, TM, Ramdoss Y, Buckler ES. 2007. TASSEL: software for association mapping of complex traits in diverse samples. Bioinformatics 223, 2633–2635.

Buhr T, Sato S, Ebrahim F, et al. 2002. Ribozyme termination of RNA transcript down-regulate seed fatty acid genes in transgenic soybean. The Plant Journal 30, 155–163.

Bus A, Hecht J, Huettel B, Reinhardt R, Stich B. 2012. High throughput polymorphism detection and genotyping in *Brassica napus* using next-generation RAD sequencing. BMC Genommics 13, 281.

Cao J, Lin W, Xie G, Ding S, Zhang X, Chen J. 2012. Evaluation on the main properties of 12 new oil palm varieties. Chinese Journal of Tropical Crops 8, 1359–1365.

Chen S, Huang Z, Dai Y, Qin S, Gao Y, Zhang L, et al. 2013. The development of 7E chromosome-specific molecular marker for Thinopyrum elongatum based on SLAF-seq technology. PLOS ONE 8, e65122.

Choi KH, Heath RJ, Rock CO. 2000. Beta-ketoacyl-acyl carrier protein synthase III (FabH) is a determimning factor in branched-chain fatty acid biosynthesis. Journal of Bacteriology 182, 365–370.

Clough SJ, Bent AF. Floral dip: a simplified method for Agrobacterium – mediated transformation of Arabidopsis thaliana. The Plant Journal 16, 735–743.

Elshire RJ, Glaubitz JC, Sun Q, Poland JA, Kawamoto K. 2011. A robust, simple genotyping-by-sequencing (GBS) approach for high diversity species. PLOS ONE 6, e19379.

Flint-Garcia SA, Thornsberry JM, Buckler ES. 2003. Structure of linkage disequilibrium in plants. Annual Review of Plant Biology 54, 354–357.

Geng X, Jiang C, Yang J, Wang L, Wu X, Wei W. 2016. Rapid identification of candidate genes for seed weight using the SLAF-seq method in *Brassica napus*. PLOS ONE 11, e0147580.

Ginestet C. 2011. Ggplot 2: elegant graphics for data analysis. J R Stat Soc A 174, 245–246.

Hardy OJ, Vekemans X. 2002. SPAGeDi: a versatile computer program to analyse spatial genetic structure at the individual or population levels. Molecular Ecology Notes 2, 618–620.

Huang X, Zhao Y, Li C, et al. 2011. Genome-wide association study of flowering time and grain yield traits in a worldwide collection of rice germplasm. Nature Genetics 44, 32–39.

Huide Huang. 2017. The industrial survey of *Elaeis guineensis* in Malaysia. World Tropical Agriculture Information 7, 7.

Hunt MC, Solaas K, Kase BF, Alexson SE. 2002. Characterization of an acyl-CoA thioesterase that functions as a major regulator of peroxisomal lipid metabolism. Journal of Biological Chemistry 277, 1128–1138.

Jeennor S, Volkaert H. 2014. Mapping of quantitative trait loci (QTLs) for oil yield using SSRs and gene-based markers in African oil palm (*Elaeis guineensis* Jacq.). Tree Genetics and Genomes 10, 1–14.

Kenny EE, Kim M, Gusev A, Lowe JK, Salit J et al. 2011. Increased power of mixed models facilitates association mapping of 10 loci for metabolic trait in an isolated population. Human Molecular Genetics 20, 827–839.

Kent WJ. 2002. BLAT-the BLAST-like alignment tool. Genome Research 12, 656–664.

Kozich JJ, Westcott SL, Baxter NT, Highlander SK, Schloss PD. 2013. Development of a dual-index sequencing strategy and curation pipeline for analyzing amplicon sequence data on the Miseq Illumina sequencing platform. Applied and Environmental Microbiology 79, 5112–5120.

Li-Beisson Y, Shorrosh B, Beisson F, Andersson MX, Arondel V, Bates PD, Franke RB, et al. 2013. Acyl-lipid metabolism. The Arabidopsis book 11, e0161.

Li H, Durbin R. 2009. Fast and accurate short read alignment with Burrows-Wheeler transform. Bioinformatics 25, 1754–1760.

Li H, Handsaker B, Wysoker A, Fennell T, Ruan J, Homer N, et al. 2009. The sequence alignment/map format and SAMtools. Bioinformatics 25, 2078–2079.

Li H, Peng Z, Yang X, Yang X, et al. 2013. Genome-wide association study dissects the genetic architecture of biosynthesis in maize kernel. Nature Genetics 45, 43–50.

McKenna A, Hanna M, Banks E, Sivachenko A, Cibulskis K, Kernytsky A, et al. 2010. The genome analysis toolkit: a Map Reduce framework for analyzing next-generation DNA sequencing data. Genome Research 20, 1297–1303.

Mekhedov S, de Ilarduya OM, Ohlrogge J. 2002. Toward a functional catalog of the plant genome a survey of genes for lipid biosynthesis. Plant Physiology 122, 389–402.

Montoya C, Lopes R, Flori A, Cros D, Cuellar T, Summe M, et al. 2013. Quantitative trait loci (QTLs) analysis of palm oil fatty composition in an interspecific pseudo-backcross from Elaeis oleifera (HBK) Cortes and oil palm (*Elaeis guineensis* Jacq.). Tree Genetics Genomes 9, 1207–1225.

Mortazavi A, Williams BA, McCue K, Schaeffer L, Wold B. 2008. Mapping and quantifying mammalian transcriptomes by RNA-seq. Nature Methods 5, 621–628.

Murray M, Thompson WF. 1980. Rapid isolation of high molecular weight plant DNA. Nucleic acids Research 8, 4321–4326.

Pritchard JK, Stephens M, Donnelly P. 2000. Inference of population structure using multilocus genotype data. Genetics 155, 945–959.

Pritchard JK, Stephens M, Rosenberg NA, Donnelly P. 2000. Association mapping in structured populations. American Journal of Human Genetics 67, 170–181.

Rajanaidu A, Kushairi A, Mohd-Pin A. 2017. Oil palm genetics resources. In: Oil palm improvement through the use of genetic resources, pp 221. Malaysia: Malaysian Plam Oil Board.

Raman H, Raman R, Coombes N, et al. 2015. Genome-wide association analyses reveal complex genetic architecture underlying natural variation for flowering time in canola. Plant Cell Environment 39, 1228–1239.

Rance KA, Mayes S, Price Z, Jack PL, Corley RHV. 2001. Quantitative trait loci for yield components in oil palm (*Elaeis guineensis* Jacq.). Theoretical and Applied Genetics. 103: 1302–1310.

Salas JJ, Ohlrogge JB. 2002. Characterization of substrate specificity of plant FatA and FatB acyl-ACP thioesterases. Archives of Biochemistry and Biophysics 403, 25.

Sasaki Y, Kozaki A, Hatano M. 1997. Link between light and fatty acid synthesis: thioredoxin-linked reductive activation of plastidic acetyl-CoA carboxylase. Proceedings of the National Academy of Science 94, 11096–11101.

Singh R, Tan SG, Panandam JM, Rahman RA, Ooi LC, Low E-TL et al. 2009. Quantitative trait loci (QTLs) analysis of palm oil fatty acid composition in an interspecific pseudo-back cross from Elaeis oleifera (HBK) Cortes and oil palm (*Elaeis guineensis* Jacq.). Tree Genetics Genomes 9: 1207–1225.

Smit AFA, Hubley R, Green P. RepeatMasker Opean-4.0. Available online at: http://www.repeatmasker.org.

Sorkheh K, Malyshevaotto LV, Wirthensohn MG, Tarkeshesfahani S, Martinezgomez. 2008. Linkage disequilibrium, genetic association mapping and gene localization in crop plants. Genetics and Molecular Biology 31, 5499–5503.

Sun X, Liu D, Zhang X, Li W, Liu H, Hong W, et al. 2013. SLAF-seq: an efficient method of large-scale de novo SNP discovery and genotyping using high-throughput sequencing. PLOS ONE 8, e58700.

Tassell CPV, Smith TPL, Matukumalli LK, Taylor JF, Schnabel RD. 2008. SNP discovery and allele frequency estimation by deep sequencing of reduced representation libraries. Nature Methods 5, 247–252.

Ting N-C, Jansen J, Nagappan J, Ishak Z, Chin CW, Tan SG, et al. 2013. Identification of QTLs associated with callogenesis and embryogenesis in oil palm using genetic linkage linkage maps improved with SSR markers. PLOS ONE 8, e53076.

Tisne S, Denis M, Cros D, Pomies V, Riou V, Syahputra I, Omore A, Durand-Gasselin D, Bouvet J-M, Cochard B. 2015. Mixed model approach for IBD-based QTL Mapping in a complex oil palm pedigree. BMC Genomics 16, 798.

Thurston M and Field D. Msafinder: Detection and characterization of microsatellite. Oxfold: CEH. Available online at: https://github.com/knirirr/Msatfinder.

Turner SD. 2014. Qqman: an R package for visualizing GWAS results using QQ and manhattan plots, BioRxiu, 005165.

Ukoskit K, Chanroj V, Bhusudsawang G, Pipatchartlearnwong K, Tangphatsomruang S, Tragoonrung S. 2014. Oil palm (*Elaeis guineensis* Jacq.) linkage map, and quantitative trait locus analysis for sex ratio and related traits. Molecular Breeding 33, 415–424.

Voelker TA, Hayes TR, Cranmer AM, et al. 1996. Genetic engineering of a quantitative trait: metabolic and genetic parameters influencing the accumulation of laurate in rapeseed. The Plant Journal 9, 229–241.

Wilson RF, Marquardt TC, Novitzky WP, et al. 2001. Metabolic mechanisms associated with alleles governing the 16:0 concentration of soybean oil. Journal of the American Oil Chemists’ Society 78, 335–340.

Zeng B, Yan H, Liu X, Zang W, Zhang A, Zhou S, Huang L, Liu J. 2017. Genome-wide association study of rust traits in orchardgrass using SLAF-seq technology. Hereditas 154, 5.

Zhang Y, Wang L, Xin H, Li D, Ma C, Ding X et al. 2013. Construction of a high-density genetic map for sesame based on large scale marker development by specific length amplified fragment (SLAF) sequencing. BMC Plant Biology 13, 141.

Zhang Z, Shang H, Shi Y, Huang L, Li J, Ge Q, et al. 2016. Construction of a high-density gentic map by specific locus amplified fragment sequencing (SLAF-seq) and its application to quantitative trait (QTL) analysis for boll weight in upland cotton (*Gossypium hirsutum*). BMC Plant Biology 16, 79.

